# Comparative Gene Essentiality across the Bacterial Domain

**DOI:** 10.1101/2020.02.28.969238

**Authors:** Daniel Shaw, Antonio Hermoso, Maria Lluch-Senar, Luis Serrano

**Affiliations:** Centre for Genomic Regulation (CRG), The Barcelona Institute of Science and Technology, Dr. Aiguader 88, Barcelona 08003, Spain; Universitat Pompeu Fabra (UPF), Barcelona 08002, Spain; ICREA, Pg. Lluís Companys 23, Barcelona 08010, Spain

## Abstract

Comparative genomics among bacteria has been used to gain insight into the minimal number of conserved genes in biological pathways. Essentiality studies have provided information regarding which genes are non-dispensable (essential, E) for cell growth. Here, we integrated studies of gene conservation, essentiality and function, performed in 47 diverse bacterial species. We showed there is a modest positive correlation between genome size and number of essential genes. Interestingly, we observed a clear shift in the functions assigned to these essential genes as genome size increases. For instance, essential genes related to transcription and translation dominate small genomes. In contrast, in large genomes functions of essential genes are related with cellular processing and metabolism. Finally, and most intriguing, we found a group of genes that while being highly conserved are also typically non-essential. This suggests that some housekeeping genes confer a significant survival benefit in nature while being non-essential *in vitro*.

## INTRODUCTION

Describing the minimal number of genes that are non-dispensable for life is one of the main challenges in biology, specifically in regard to the rational engineering of a living system. In recent years, theoretical and experimental studies have addressed the question of what constitutes a minimal genome, mainly using prokaryotes as model organisms (1–5). While there are many fundamental processes common to all un-cellular life (6–9), bacteria inhabit almost all known niches on the planet, from intracellular parasites to the Atacama desert (10–12). As such, establishing a common metabolome under such vast differences in nutrient availability and composition is unlikely (1,7).

Studies investigating the conservation of genes by genome comparison in prokaryotes have come out with many different numbers. Brown et al. found 23 genes conserved among 45 genomes (13), Harris et al. found 80 genes common to 34 genomes (14), Koonin found 63 conserved genes from approximately 100 genomes (15). When only 7 minimal endosymbionts and parasites were compared the value rises to 156 (16). In all cases, the number of conserved metabolic and structural genes rises comparatively to the decrease in diversity of the pool of organisms studied, as they are either closer evolutionarily, or have similarities in lifestyle or niche (endosymbionts and parasites). Charlebois and Doolittle (17) analysed 130 bacterial genomes and found around 62 conserved genes. More recently, analysis of 2000 fully sequenced bacterial genomes identified 73 additional marker genes that are each present within at least 90% of all prokaryotes (18).

On the other hand, experimental genome reduction coupled with transposon mutagenesis has resulted in the creation of a minimal bacterial genome consisting of 473 genes, of which 149 have unknown functionality (2). Essentiality studies in naturally occurring genome reduced bacteria such as *Mycoplasma pneumoniae* and *Mycoplasma agalactiae* show that 301 and 303 protein coding genes are essential, respectively (4,19). This disparity in numbers points to the fact that conservation of functionality (i.e. the presence of a chaperone) is more important than conservation of a specific gene. Genetic redundancy allows bacteria to survive the loss or disruption of a specific gene, either through duplication (20), or via the presence of a protein that exhibits a moonlighting function able to recover the lost phenotype (21). Finally, the fact that in some cases two different genes can perform the same function, like in the case of some aa tRNA synthetases (15) also explains the lower number of universally conserved genes (14–17). Comparative analysis of essentiality across 23 bacterial species has shown that essential (E) genes are more evolutionarily conserved than non-essential (NE) genes (22).

In recent years, multiple studies have addressed the essentiality of genes in a specific bacterium; mainly via transposon mutagenesis studies (see suppl. Table S1). In these studies, the resolution depends on the saturation of insertions. High saturation allows for proper classification of genes of different lengths as essential, non-essential or fitness (F) (23,24). E genes cannot be disrupted or removed without killing the bacteria, and are by definition non-dispensable for cell survival. NE genes can be deleted without initiating a lethal phenotype (1,25). The third category, F genes, also known as quasi-essential genes (2), are those that can be deleted from the genome but incur a significant cost to the viability of the organism (4). Other methods like site directed mutagenesis could also be used to classify gene essentiality, but lack the power offered by deep sequencing and transposon insertion (26). When doing transposon essentiality studies it is important to note that essentiality of some genes will depend on the experimental conditions (biological dependent essentiality) and on the number of passages (experimental dependent essentiality).

As complexity within the genome increases, redundancies in essential functions in principle should allow for fewer genes to be labelled as essential (27). However, it is possible that as genome size increases, and more modules and functions are added, there could be a proportional increase in the number of critical nodes. This in turn could lead to an increase in the number of E genes. This relationship has been explored analogously by Basler et al, showing that metabolic networks of different sizes demonstrate the same pattern with regard to the number of “driver reactions” (28). As the complexity of the system increases, more points of failure become present, and certain NE functions become integrated into new essential circuits, changing their original essentialities.

Essentiality studies combined with the presence of high numbers of sequenced genomes allows us to answer the following questions: a) What is the minimum number of E genes in all species studied? b) Are the highly conserved genes always E or could we find conserved NE genes? c) Does increasing genome complexity imply a higher number of E genes due to increased critical node number or epistatic interactions? d) Does the function of the cell’s E genes change depending on genome size? In other words, large genomes will have more essential genes in a given functional category, while small genomes will have a different composition?

Here, we tried to answer the above questions. We showed that both the size and composition of the essential genome of a bacteria changes as the complexity of the genome increases. In addition, we showed that universally conserved genes have a strong bias towards functions relating to transcription, translation and DNA replication & repair. However, the essentiality of these genes is not strongly conserved, and universally E genes are rare. We also identify a subset of genes that are highly conserved, yet rarely essential. We show how the essentiality of each COG category changes with genome size, and that while there are few individual examples of genes becoming more or less essential as genome size changes, global trends relating to COG category essentiality do emerge.

Finally, we show that there is a non-linear correlation between the level of gene conservation and its essentiality. Genes that are present in less than 30 species become less essential as they are included in more species, yet once a gene is present in more than 30 species, it becomes the likelihood of it being E increases.

## MATERIAL AND METHODS

### Database creation

A PubMed search was initiated using the search terms “Essential genes” and “Bacteria”. 107 separate entries were found listing the essential genes of a specific bacterial species, corresponding to 84 different bacterial species across 68 papers. These results were filtered to allow for a standardised comparison across the different data sets. As such, the studies were filtered via three categories: i) The authors must have used mini-transposons do disrupt the genome; ii) Insertions and thus gene essentiality were determined via a Next Generation Sequencing methodology; and iii) The paper must provide a list of genes deemed essential for the organism being studied. In addition, studies done in *Escherichia coli* and *Bacillus subtilis* by using systematic knockouts of every gene within the genome, were deemed to be representative for this study and were added. 47 entries matched the inclusion criteria and were included in Suppl. Table 1, spanning 8 phyla (Suppl. Figure 1 & Suppl. Table 2).

For each species strain, the relevant genome assembly was downloaded to build up a relational database. Wherever possible, RefSeq datasets were preferred. GenBank datasets where chosen if RefSeq ones did not exist for that strain. FASTA sequence files were gathered from each dataset jointly with extracted GenBank-formatted information for each entry extracted with NCBI Entrez Direct command-line tools (29). If .gff annotation files were available, they were also used with the previous other files in order to collect different synonyms for each entry identifier. All this information was stored in a specific table in the relational database.

The datasets above were parsed to remove all pseudogenes and RNAs from the analysis, ensuring that only protein coding sequences were analysed. For each species, the list of E genes was extracted from the appropriate paper. The list of E genes was cross-referenced to the genes within the annotations for that species, ascribing an essentiality status of E or NE to each gene. Since in many studies they don’t distinguish between F and NE genes, we included F genes into the NE category. Any cases where the two lists did not match automatically were manually annotated, using BLAST to identify homologs.

COG assignations were extracted from the available files. All genes with a RefSeq ID or GenBank ID that contained a GOG assignation was ascribed with a corresponding COG category. Any gene without a COG category assigned, but having otherwise a homolog with an assigned COG one, was also ascribed with the functional annotation of the homolog. Any cluster that contained a “S” annotation, denoting “Unknown Function” as well as a different annotation had the “S” annotation removed, as the function that had been ascribed to one member of the cluster was ascribed to them all, thus the “S” COG became redundant. COGs were grouped into four Super-COGs for general analysis, which are shown in Suppl. Table 3.

All previous assignations and annotations were also stored in separate relational tables that could be interlinked to the original table containing entries IDs. The same approach was followed for subsequent analyses data, so it became easier to design queries for retrieving aggregated total numbers or relevant lists, such as the ones presented in the supplemental material.

### Gene homologue clustering and assignment

Genes were clustered into groups of homologs using ProteinOrtho. This tool generates clusters containing protein sequences grouped according to their mutual similarity by using the results of successive BLASTP runs. For our case, we used an E-value parameter of 1e-05, a similarity of “0.95”, an identity of “25” and a coverage of “50” (full list of parameters can be found in Suppl. Table 4). Duplicated genes in a single species were filtered out so only one record remained, and fusion proteins were treated as a single protein or two proteins, depending on their RefSeq ID. In total, 63923 unique clusters of homologs were established.

To try and keep everything as standardised as possible, we attempted to use the same genome annotation format throughout and cluster genes with homologs. This could allow annotated species to infer and double check annotations from less well defined species. In this spirit, we decided to focus on ORFs encoding for proteins, excluding functional or non coding RNAs from the analysis. Within the issue of standardisation, we encountered a wide variety in the quality and completeness of the annotations used. While we tried to standardise all of the genes from each organism in our database to be linked to a RefSeq ID, this was not always possible. As such, GenBank IDs, and rarely other IDs, were also used to ensure that there were no gaps in our records for each species. On top of this, matching the IDs given for the essential genes to our database for the species often non-trivial. This was due in part to the variety of reporting methodologies used by each author, and in part to a lack of synchronisation between the genome annotations and the list of E genes provided.

As a point of clarity, we use the phase ‘homolog’ to define genes that cluster together using our ProteinOrtho search, but are found in different organisms. ‘Paralog’ is used to define genes that perform identical functions but do not appear in the same cluster, due to divergent sequence identity.

For the list of E genes from each species, an essentiality indicator was established from each paper. For example, the list of essential genes provided for *Synechoccus elongatus* contained the RefSeq ID for each E gene (30), so matching this against our database was easy to do automatically, and allowed us to easily annotate which genes in the database relating to *S. elongatus* were E. Most lists gave a locus ID that matched to the genome annotation, such as those given for *Streptococcus pyogenese* (31), and some such provided genomic loci for each gene, along with other identifiers, such as *Herbaspirillum seropedicae* (32). In general these were fairly simple to match, however there were ambiguous cases. For those lists that contained other identifying information, such as genetic loci, resolving these miss-matches was much easier. Some papers provided only common genes names, like the list provided for *Bacillus subtilis* (33). This was most problematic, as many genes have multiple common names, such as the Ribosomal RNA small subunit methyltransferase A being referred to as either *rsmA* or *ksgA* interchangeably (34).

Due to the large amount of variation in the input data, this meant that many E genes had to be identified manually, as they did not match directly to the database. In such cases, we generally ran BLASTP to identify if there were any obvious homologs in our dataset already if the sequence of the protein was known. If this was not the case, descriptions of the proteins function were used to see if it could be mapped to a protein in that species from the database.

These efforts were hampered further by a mix of miss-annotation of E genes and changes to the genome annotation files after the papers had been published. For example, in the list of essential genes provided for *Bacillus thuringiensis*, there are five typos in the gene names (35). The essential Asparginase in this organism is labelled with the incomplete locus tag “BMB171_C1”. Looking into the genome, *B. thuringiensis* contains two Asparginases, BMB171_C2086 and BMB171_C1329. As no other information on the gene is provided, it was annotated as BMB171_C1329 on the basis of the partial locus tag. Other errors, such as underscores (_) being replaced with hyphens (-) were less ambiguous to correct, but still required manual curation.

Other times, the genome annotation had been updated or modified since the paper relying on it had been published. A good example of this can be found in the annotations of *M. tuberculosis*, where the genes Rv3021c and Rv1784 are designated as E (36). However, in the genome annotation we downloaded of *M. tuberculosis*, Rv3021c was annotated as a pseudogene and Rv1784 no longer existed, as it had been determined that it was actually a part of Rv1783, not a unique protein itself. As such, both annotations were discarded, as we discarded all pseudogenes from the analysis (on the basis that they are not genes) and Rv1783 was already annotated as E.

All of our clustering techniques were based on protein identity, grouping genes with a similar protein structure with each other and assuming homology of structure equals homology of function. However, this ignores the fact that there is often more than one gene responsible for the same phenotype in different bacteria (37). As a standard, we used the genes in *M. pneumoniae* as the basis for the initial clustering, as they are well described and annotated (38). However, just because a gene performs a specific function in *M. pneumoniae*, does not mean all other bacteria that contain that function will contain that specific gene. While the *M. pneumoniae* genes were used as a base, we clustered the entire database by homology, so all gene groups that are homologs are successfully clustered. The issue comes when querying the database about information for a specific gene.

A clear example of this was found in the genes coding for the proline tRNA synthetase. According to Charlebois & Doolittle (17) and Koonin *et al*., (15), we should find the proline tRNA synthetase in every species. However, the copy of the gene found in *M. pneumoniae* only had homologs in eight other species. We therefore had to manually query every species to see if it contained an annotated proline tRNA synthetase, and we found that there were three distinct paralogs of the gene that were generally split along phylogenetic line, though two species contain two different versions of the gene (*B. thuringiensis* & *B. phytofirmans*). Notably, while many species lack an asparagine and glutamine tRNA synthetases, they contain the *gatA/gatB* system, which amidates Asp and Glu loaded on Asn-tRNA and Gln-tRNA (39). Having different classes of tRNA synthases has been well documented (40), and serves as a reminder that evolution has allowed for multiple different paths to the same outcome.

Another problem happens when a small gene is fused to another, is elongated or has a long insertion. An example is the *M. pneumoniae* copy of *nusG*. When we searched the database, we found no other species that contained the gene. The reason being that it the gene is much longer in *M. pneumoniae* and has an insertion in the most conserved region (See Suppl. Figure 2). In those cases, manual sequence comparisons should be performed. This issue of search results being biased due to the expectation of preserved homologs gives further credence to the secondary search strategy employed by Charlebois & Doolittle (17), where they ascribed a function and gene name to each homolog cluster they collated. Searching databases via protein sequence or a RefSeq ID alone will inevitably bias the results towards that specific homolog, ignoring important paralogs. While there is a huge variance in the gene name annotations given, the ability to search via function instead of a specific gene would be a huge help in identifying common features among multiple species.

### Clustering essentiality by genome size

All clusters of homologs containing an essential gene were isolated and divided by COG category. They were then plotted against all bacterial species in order of genome size to form a heat-map. A green colour indicates the gene was present in that organism but was NE, a red colour indicates the gene was present and E, and a white colour indicated the gene was not present.

## RESULTS

### Universally conserved genes and their functions

By performing a bibliographic search, we collected data of essentiality from 107 entries, which were selected as indicated in Methods. After cleaning by the methodology used and completeness of the study, a total of 47 species, represented by 8 phyla, were selected to generate the essentiality database that was used in posterior analysis (Suppl. Tables 1, 2 and Suppl. Figure 1). This database contained 191341 genes that can be further split into 63923 clusters of homologs (As defined in Methods). Of the 63923 clusters of homologs in the database, we could assign COG categories to 15823 (25%) of them. However, these 15823 clusters with a COG annotation account for 109549 of the 191341 (57%) of the genes in the database. The composition of the database via COG categories is shown in Suppl. Table 5.

Study of gene conservation revealed that only 92 out of 63923 clusters were universally conserved (Table 1), and allowing for one species missing a gene (to consider error of annotation or sequence search) the number raised to 127. We compared our list of universally conserved genes with studies previously performed by other groups (See Suppl. Table 6) and found that all genes previously described by Charlebois and Doolittle (17) as universally conserved were also found within our dataset of 140 genes. However, only 70% of the genes identified by Charlebois and Doolittle had an identical homolog shared by all species. The remaining 30% of genes were still present in all species, but via paralogs.

**Table 1.**
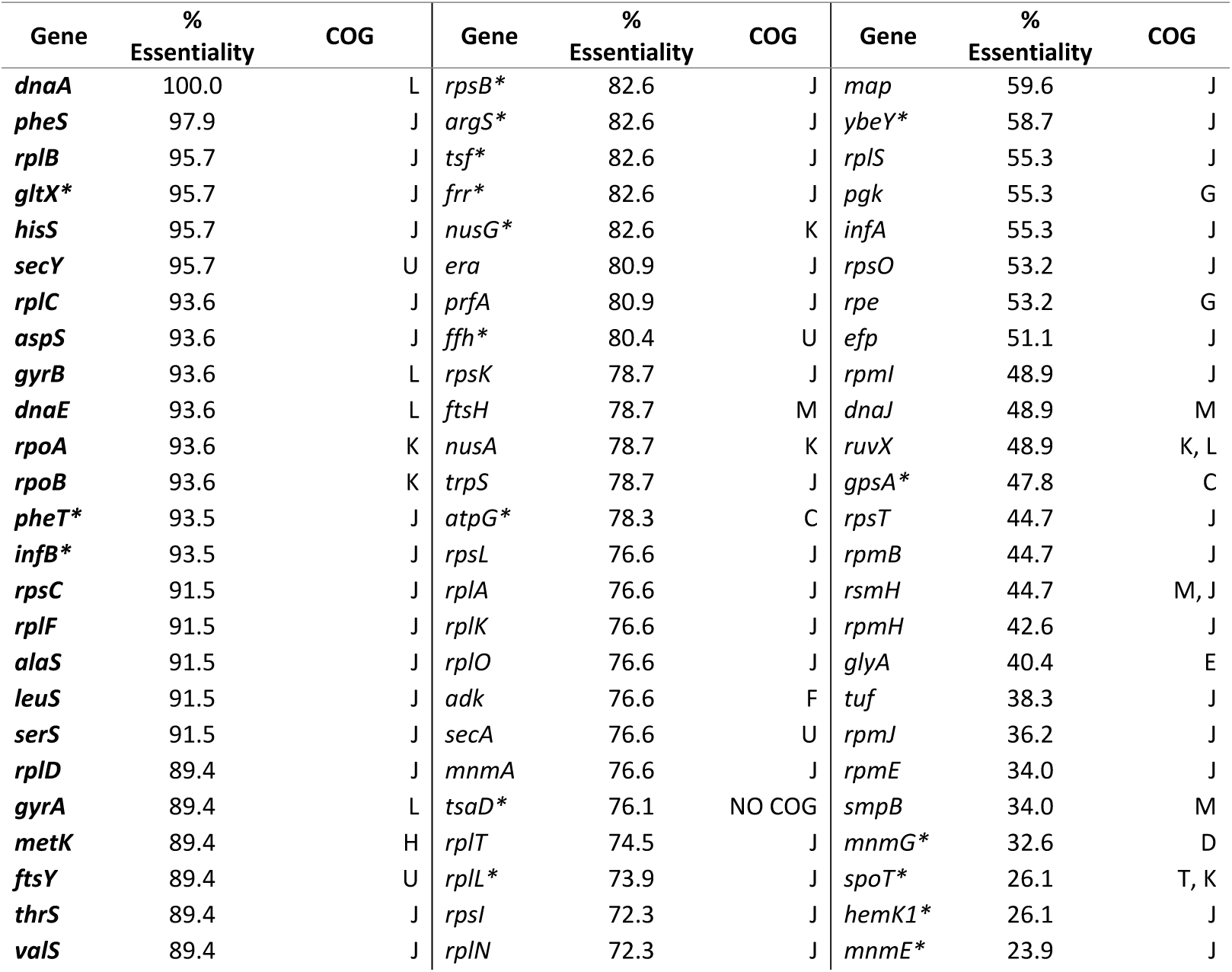
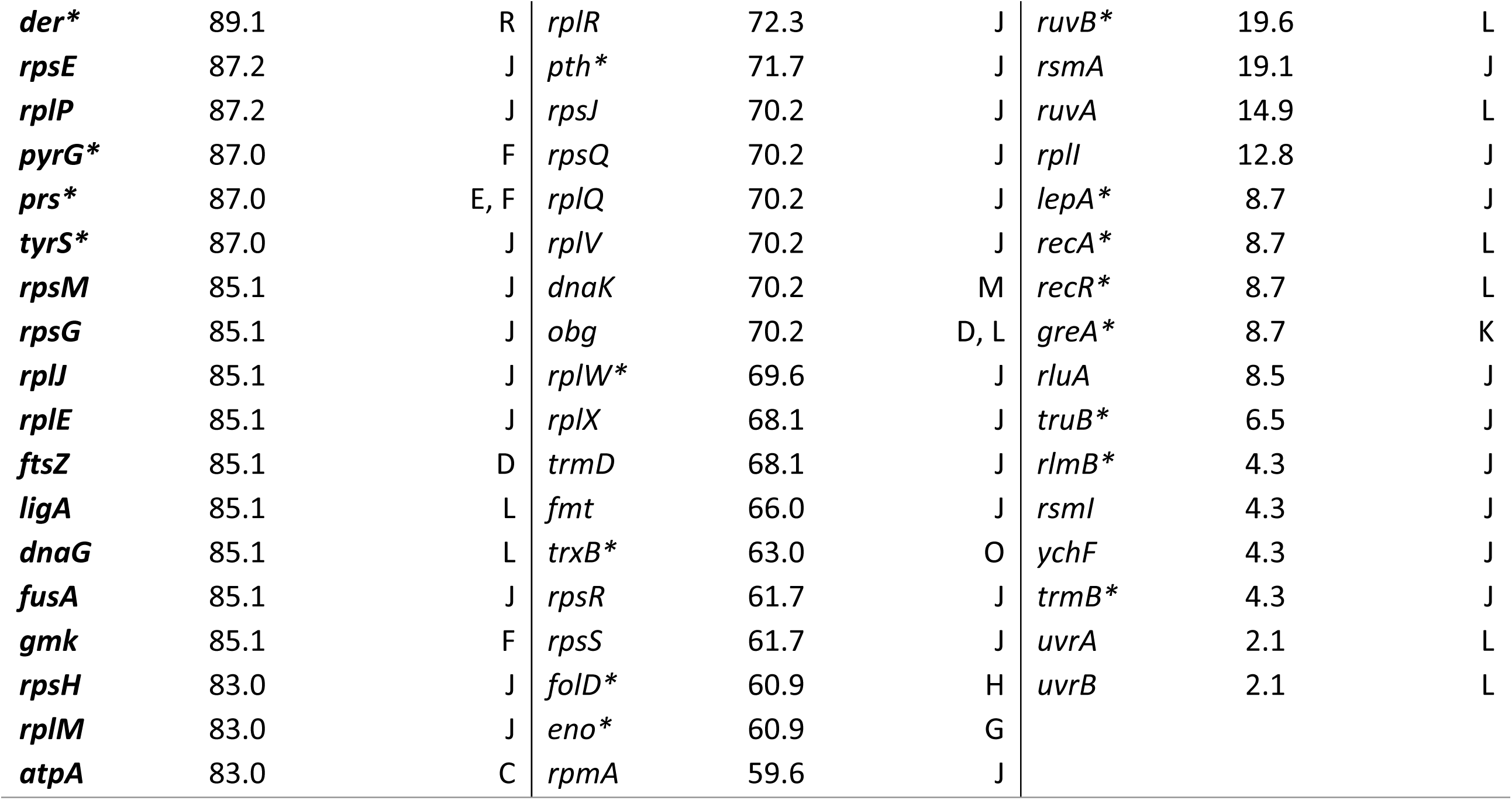
List of 92 genes universally conserved in our dataset, as well as the 35 genes missing in only a single species indicated with a (*), showing standard gene name, the percentage of species in the database that the gene is annotated as essential in, and COG category(s).

Of the list of near universal genes generated by Lan et al., (18), 60% of them had a homolog in all species in our dataset, 30% were universally conserved but with multiple paralogs and 10% were not universally conserved. In the following text, we will use the term universally conserved genes for the 127 genes we have found in our database that are found in all species, or missing in just one.

The study revealed that the majority of those universally conserved genes generally fall into one of four functional categories: i) Ribosomal proteins, or those associated to them (38%). ii) Proteins involved in signal transduction and post-translational modifications (6%). iii) Proteins that interact with the synthesis, replication or repair of DNA and RNA (24%), and iv) tRNA synthetases, and proteins associated with them (17%) (Figure 1). When looking for functionality and not gene similarity we find that all species encode for 18 tRNA synthetases and either they have Asn and Gln-tRNA synthetases or they have the GatA/GatB system capable of amidating Asp and Glu loaded on Asn and Gln tRNA (39).

**Figure 1.**
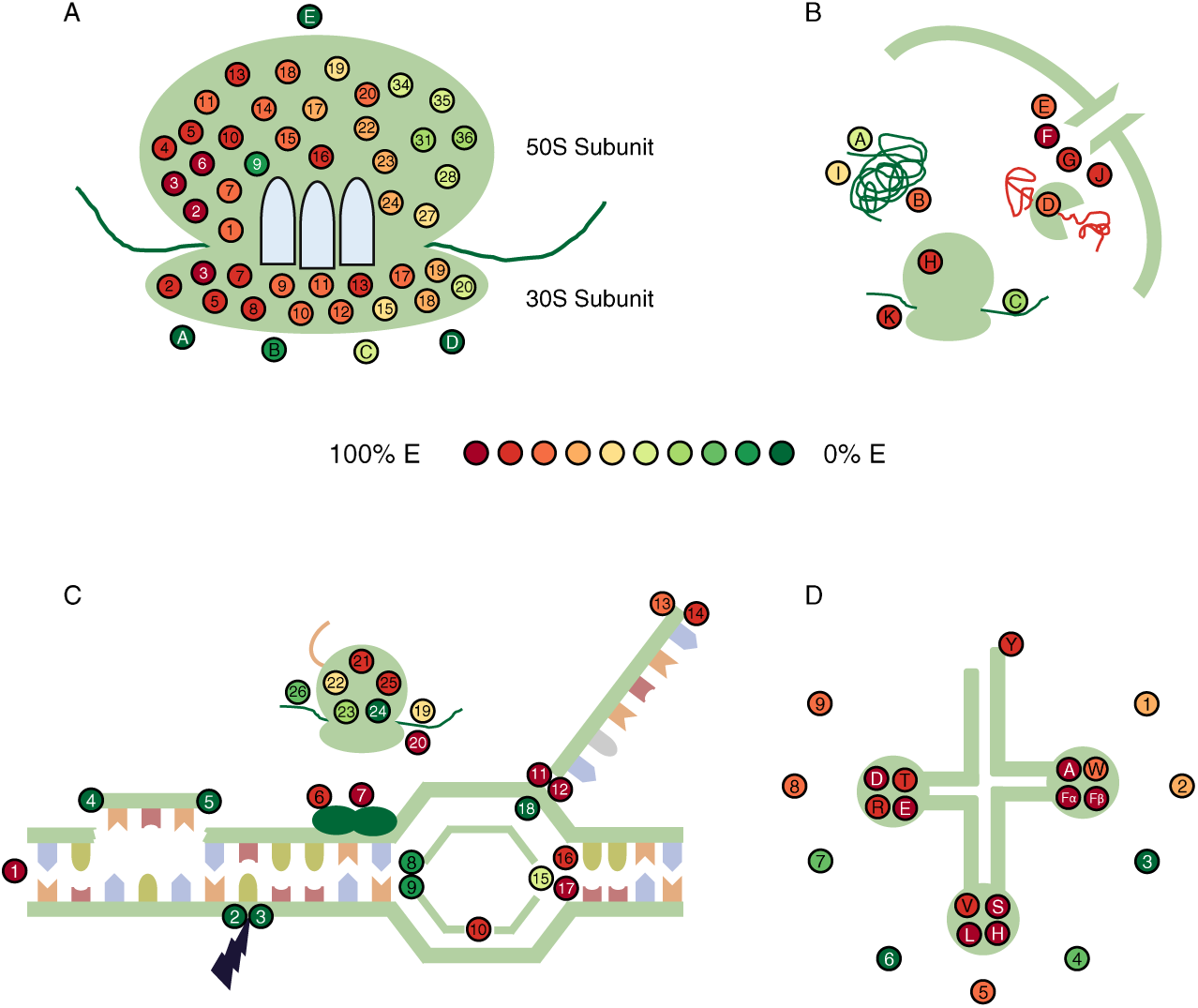
Essentiality profiles of the conserved genes, grouped by functional categories. Gene essentiality is indicated by the panel in the centre. [A] Ribosomal proteins. Number refers to that Subunit’s specificity, i.e. gene 3 in the 50S denotes ribosomal protein 50S L3. A- 23S rRNA gunosine methyltransferase *rlmB*. B- Ribosomal RNA small subunit methyltransferase A. C- Ribosomal RNA small subunit methyltransferase H. D- Ribosomal RNA small subunit methyltransferase I. E – Ribosome-binding ATPase *ychF*. [B] Genes involved in signal transduction and post-translational modification. A- Chaperone *dnaJ*, B- chaperone *dnaK*, C- ssrA-binding protein, D- ATP-dependant zinc metalloprotease *ftsH*, E – protein translocase subunit *secA*, F - protein translocase subunit *secY*, G – Signal recognition particle receptor *ftsY*, H – Peptide chain release factor 1. I- Methionine aminopeptidase map. J- Signal recognition particle *ffh*. K- Ribosome-recycling factor *frr*. [C] Genes involved in DNA & RNA synthesis, replication and repair. 1– Chromosomal replicaton initiator protein *dnaA*. 2 – UvrABC system protein A. 3 – UvrABC system protein B. 4- Recombination protein *recA*. 5- Recombination protein *recR*. 6- DNA primase *dnaG*. 7- DNA polymerase III subunit alpha. 8- Holliday junction ATP-dependant DNA helicase *ruvA*. 9- Holliday junction ATP-dependant DNA helicase *ruvB*. 10- DNA ligase *ligA*. 11- DNA-directed RNA polymerase subunit alpha. 12- DNA-directed RNA polymerase subunit beta. 13- Transcription termination/antitermination protein *nusA*. 14- Transcription termination/antitermination protein *nusG*. 15- Holliday junction resolvase *ruvX*. 16- DNA gyrase subunit A. 17- DNA gyrase subunit B. 18- Transcription elongation factor *greA*. 19- Translation initiation factor IF-13 20- Translation initiation factor IF-2. 21- Elongation factor G. 22- Elongation factor P. 23- Elongation factor Tu. 24- Elongation factor 4. 25- Elongation factor Ts. 26- Release factor glutamine methyltransferase. [D] tRNA synthetases, indicated via common 1-letter abbreviation. 1- Methionyl-tRNA formyltransferase *fmt*. 2- tRNA guanine methyltransferase *trmD*. 3- tRNA guanine methyltransferase *trmB*. 4- tRNA modification GTPase *mnmE*. 5- tRNA N6-adenosine threonylcarbamoyltransferase *tsaD*. 6- tRNA pseudouridine synthase B. 7- tRNA uridine 5- carboxymethylaminomethyl modification enzyme *mnmG*. 8- tRNA-specific 2-thiouridylase *mnmA*. 9- Peptidyl-tRNA hydrolase.

### Gene essentiality analysis of universal genes

We then examined the gene essentiality of all the genes in our database. Interestingly, only the *dnaA* gene is annotated as essential in all 47 species. This could be due to the fact that we considered F genes as NE, but in transposon studies sometimes E genes can be classified as F. For example, insertions found at the beginning or at the end of a gene could have no overall effect on the protein function, or the protein could contain a NE extension full of insertions. As a result, a gene could be classified as F or even NE instead of E (see Suppl. Figure 2).

To account for these cases, we established a threshold to define universally essential genes. For this purpose, we considered that the universally conserved alpha and beta subunits of the RNA polymerase and all non-duplicated tRNA synthetases should be E. Thus, we set a threshold for universal essentiality at 89%, and applying this threshold to all the list of candidate genes, we could identify 22 universally essential genes in our analysis (See Table 1). There are also many genes that are highly conserved yet rarely, if ever, E. We looked for genes that were found in ≥90% of species, but whose cluster had an essentiality of <10%. 57 clusters of homologs were returned, of which 20 were NE in all species (Figure 2 & Suppl. Table 7). Similarly with the universally conserved proteins, this list is dominated by genes relating to the Information storage & Processing. However, while that most highly represented class of genes (66.3%) in the universally conserved list related to translation, ribosome structure and biogenesis (COG category J), in the rarely E genes relating to DNA replication, recombination and repair were the most common, followed by translation, ribosome structure and biogenesis (33.3% and 26.3% respectively). There was also a large number of genes with only general or no function ascribed to them (COG categories R and S, 19.3% combined).

**Figure 2.**
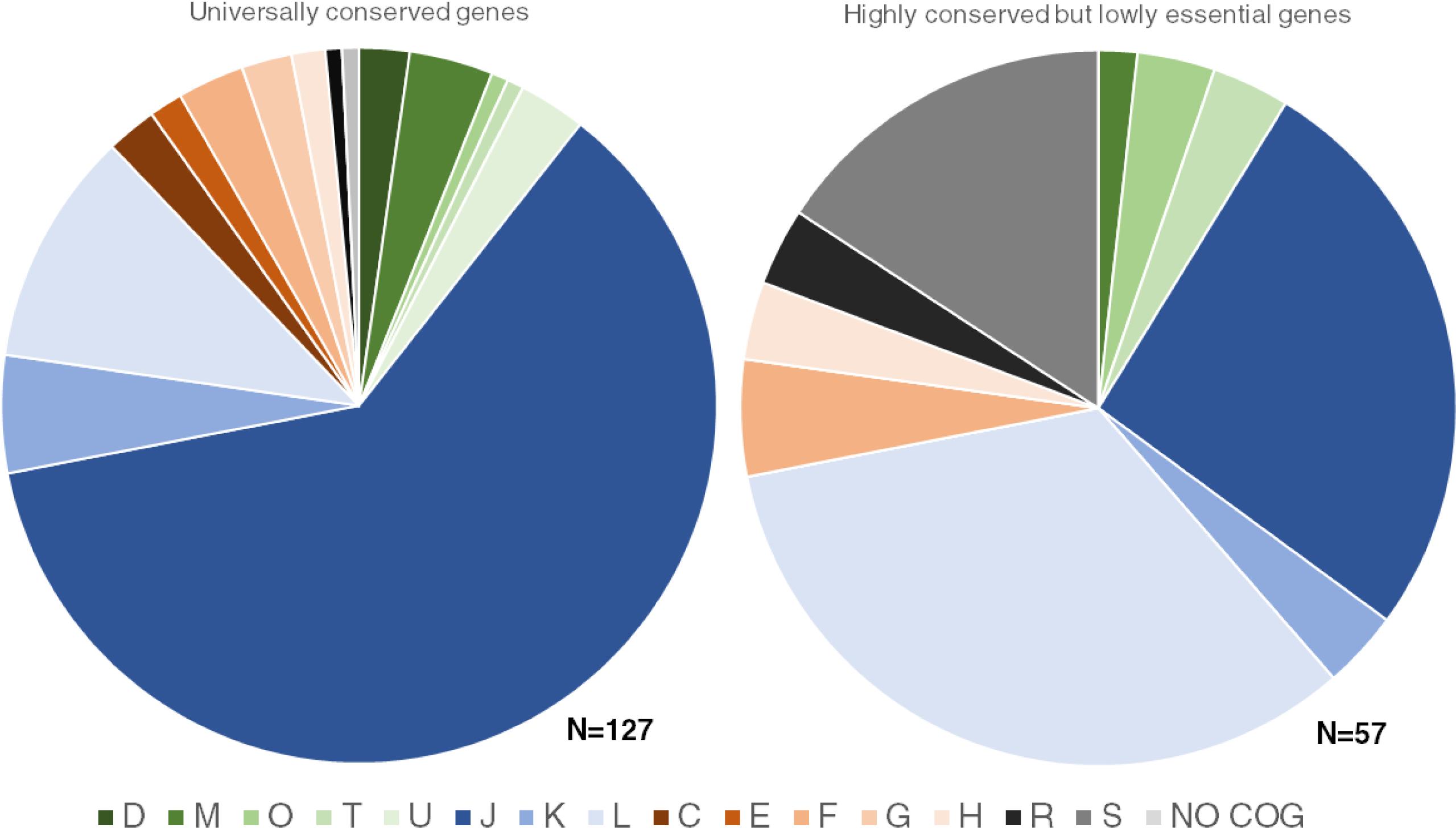
Composition of the highly conserved but lowly essential genes and the universally conserved genes via COG category. D - Cell cycle control, cell division, chromosome partitioning. M - Cell wall/membrane/envelope biogenesis. O - Post-translational modification, protein turnover, and chaperones. T - Signal transduction mechanisms. U - Intracellular trafficking, secretion, and vesicular transport. J - Translation, ribosomal structure and biogenesis. K – Transcription. L - Replication, recombination and repair. C - Energy production and conversion. E - Amino acid transport and metabolism. F - Nucleotide transport and metabolism. G - Carbohydrate transport and metabolism. H - Coenzyme transport and metabolism. R - General function prediction only. S – Function unknown.

### Genome complexity and essentiality

To evaluate the relationship between genome complexity and essentiality, we studied the correlation between the number of E genes and the genome size (total number of genes) in our database. As shown in Figure 3A, our dataset contains species with a large variety of genome sizes. The value of the Pearson’s Product-Moment correlation was between genome size and number of E genes was 0.28 (*P=0*.*055)*. This indicated that the number of E genes could be higher in more complex genomes.

**Figure 3.**
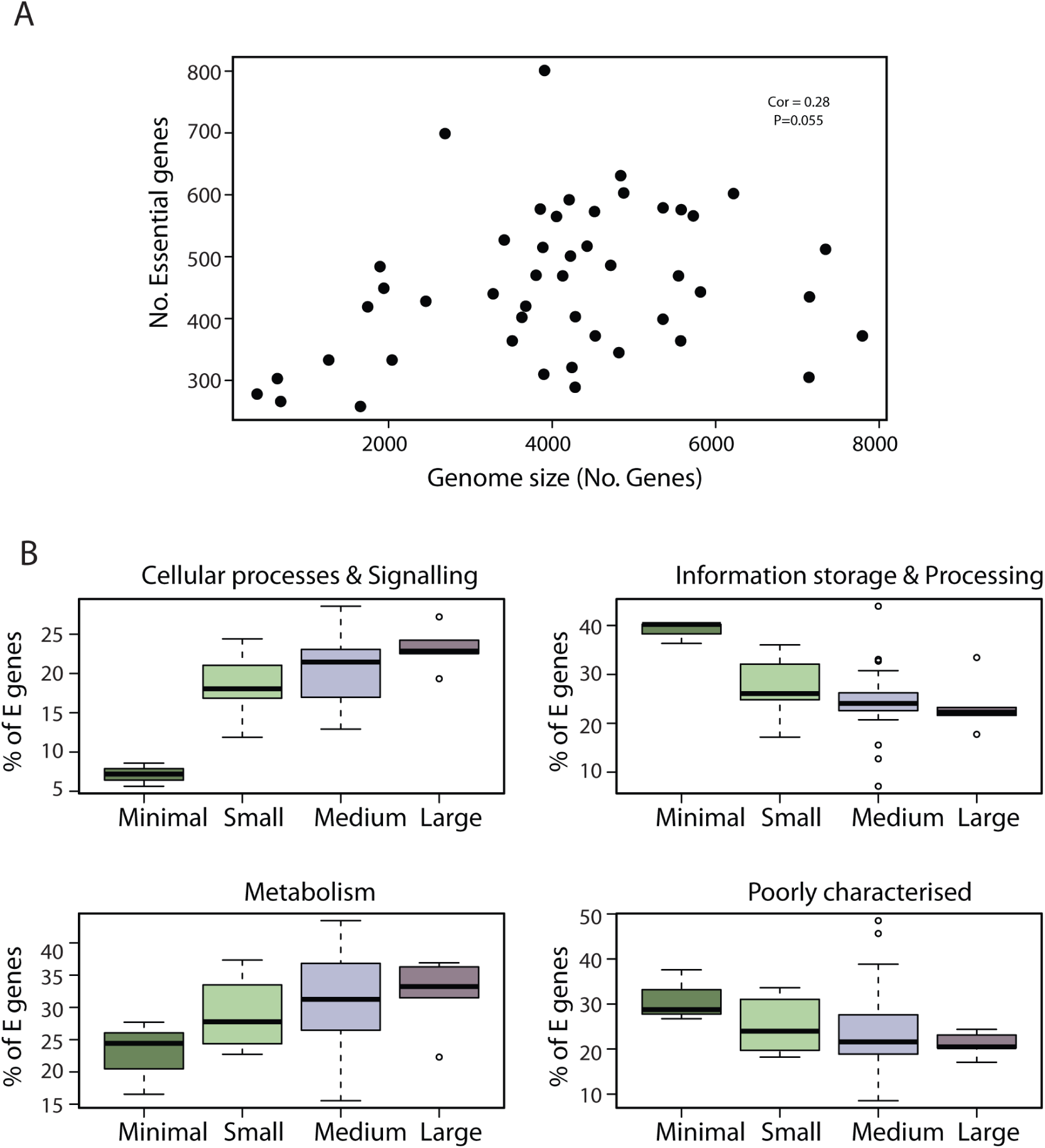
Relationship between genome size and essential genes. A - Number of essential genes vs total genome size of the 47 species studied. B - Composition of the essential genome via Super COG classes as a percentage of the total essential genome, stratified by genome size.

To gain insight into the relationship between essentiality and complexity in functional pathways, we studied if the composition of a cells’ essential genome changed depending on the genome size. First, the 47 selected bacterial were classified into four categories based on their genome sizes; minimal genomes (<1000 genes), small genomes (1000-3000 genes), medium genomes (3000-6000 genes) and large genomes (>6000 genes). Secondly, the COG categories were also grouped into four super categories; ‘Cellular processes and signalling’, ‘Information storage and processing’, ‘Metabolism’, and ‘Poorly characterised’ (See Suppl. Table 3). Then, we studied the relation between genome size and essentiality in different functional groups (Figure 3B).

For genes involved in ‘Metabolism’ and ‘Cellular processes and signalling’, there is a generally positive correlation between the increase in genome size and their percentage contribution to the essential genome. However, for genes related to ‘Information Storage & Processing’ and ‘Poorly characterised’ the number of essential genes per super COG is relatively stable, independent of genome size (Suppl. Table 8). For each of the COG categories within the super COG, the essentiality was plotted against the genome size category to see if the individual COG categories aligned with the change in the super COG, and the pattern of the individual COG categories tends to follow that of their super COG category (Suppl. Figures 3-10).

Looking at the overview of the distribution of individual essential genes across COG category and genome size (Supp. Figures S10-13), we see few clear examples of genes become more or less E as the complexity of the genome increases. However, more trends appear at the COG level. For example, E genes for cell motility (COG category N) and signal transduction mechanisms (COG category T) tend to cluster in the larger organisms, yet are much rarer in smaller ones.

While the relationship between genome complexity and essentiality is shown in Figure 3 on a global level, Figure 4 shows the relative contributions of each individual COG category has on the total essential genome in minimal genomes vs the largest genomes. The pie charts show both the size of the average essential genome, with the large genomes having on average 434 ± 101.3 (1SD) essential genes compared to the minimal bacteria’s 311 ± 22 (1SD), and their overall makeup in regards to the super-COGs. The bias towards translational and ribosomal genes in the minimal bacteria is readily apparent, along with the lack of essentiality in the cellular process and housekeeping genes.

**Figure 4.**
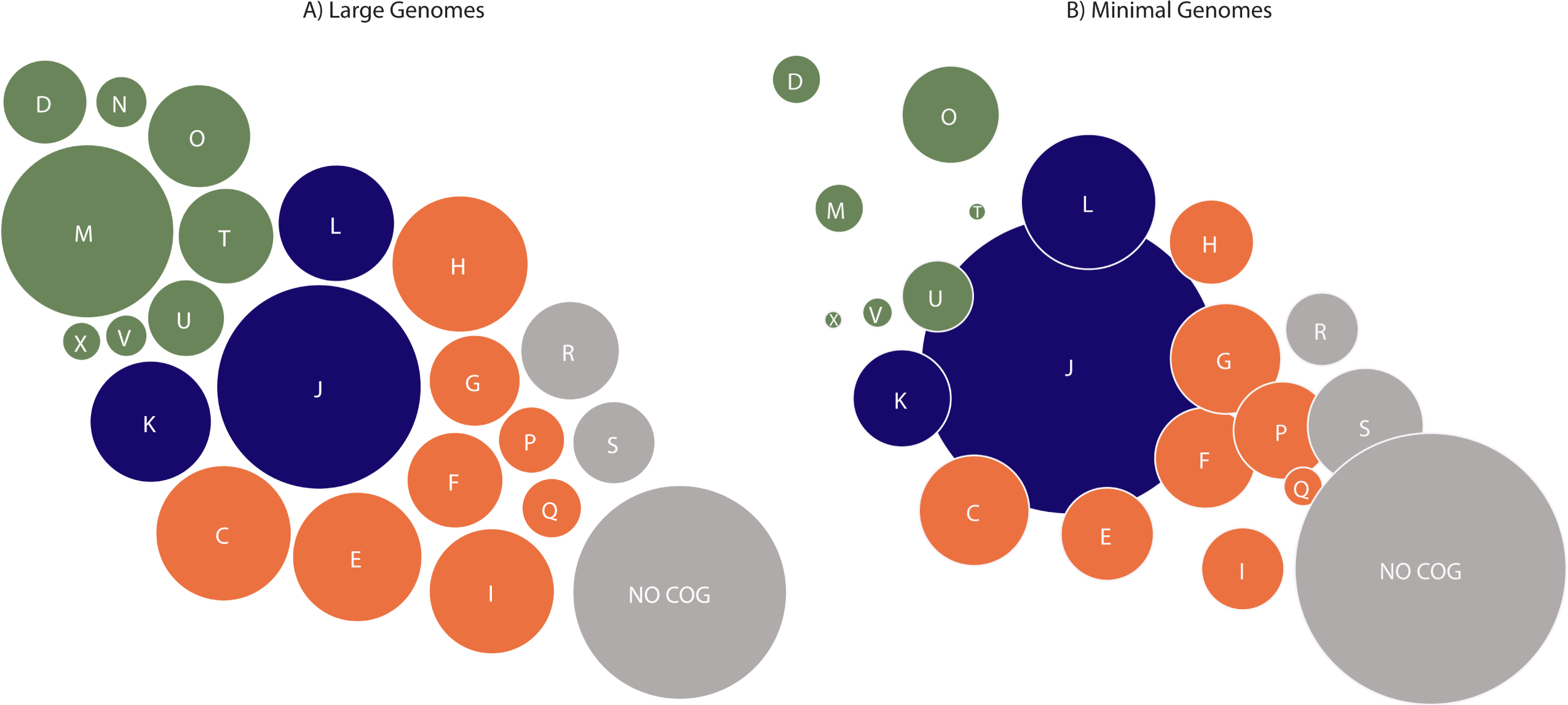
Bubble plot showing the average composition of the essential genomes of the minimal vs large bacteria by COG category, as a percentage of their total essential genes. Circles represent the relative abundance of each COG category. Green circles denote COG categories belonging to the Cellular processes & Signalling Super-COG, blue circles represent COG categories belonging to the Information storage & Processing Super-COG, orange circles represent COG categories belonging to the Metabolism Super-COG, and grey circles represent COG categories belonging to the Poorly characterised Super-COG. D - Cell cycle control, cell division, chromosome partitioning. M - Cell wall/membrane/envelope biogenesis. N - Cell motility. O - Post-translational modification, protein turnover, and chaperones. T - Signal transduction mechanisms. U - Intracellular trafficking, secretion, and vesicular transport. V – Defence mechanisms. X – Mobilome. J - Translation, ribosomal structure and biogenesis. K – Transcription. L - Replication, recombination and repair. C - Energy production and conversion. E - Amino acid transport and metabolism. F - Nucleotide transport and metabolism. G - Carbohydrate transport and metabolism. H - Coenzyme transport and metabolism. I – Lipid metabolism. P – Inorganic ion transport and metabolism. Q – Secondary metabolite biosynthesis, transport and catabolism. R - General function prediction only. S – Function unknown.

*B. thuringiensis* appeared to be an outlier from the rest of the species analysed in regard to pattern of E genes, specifically within the Information Storage & Processing Super-COG. As the eighth largest organism by genome size, its bias towards non-essentiality stands out clearly in COG category J, and to a lesser extent in categories K and L (See Suppl. Figure 12). This deviation from the standard pattern of essentiality can be explained by the fact there are no ribosomal proteins classified as essential, according to the paper investigating *B. thuringiensis* (35). This lack of ribosomal genes is not mentioned in the original paper, and analysis of the organism at the genome level does not indicate multiple copies of the proteins to provide redundancy. Therefore, either they were excluded deliberately or lost to experimental noise.

### Gene essentiality correlates in a non-linear way with gene conservation

Of the 191341 genes in our database, 21796 are classified as essential in at least one species. Theses E genes can be divided among 5510 clusters of homologous genes. However, the essentiality of a gene in one organism does not infer essentiality in all others, and there are 42471 separate NE genes that have a homolog that is E in another species. On average, each cluster of homologous genes that contains an E gene is represented in 10.6 species (SD ± 12.8), and the genes in the cluster are essential 51.8% (SD ± 37%) of the time. As previously described in Luo et al., (22), the pattern of a clusters essentiality changes based on how well conserved it is, as shown in Figure 5. As the number of species a cluster is found in increases, the chance of that gene being essential decreases. However, once a cluster is found in more than 30 species, its essentiality increases, thus the chance of it being an essential gene in more organisms begins to rise.

**Figure 5.**
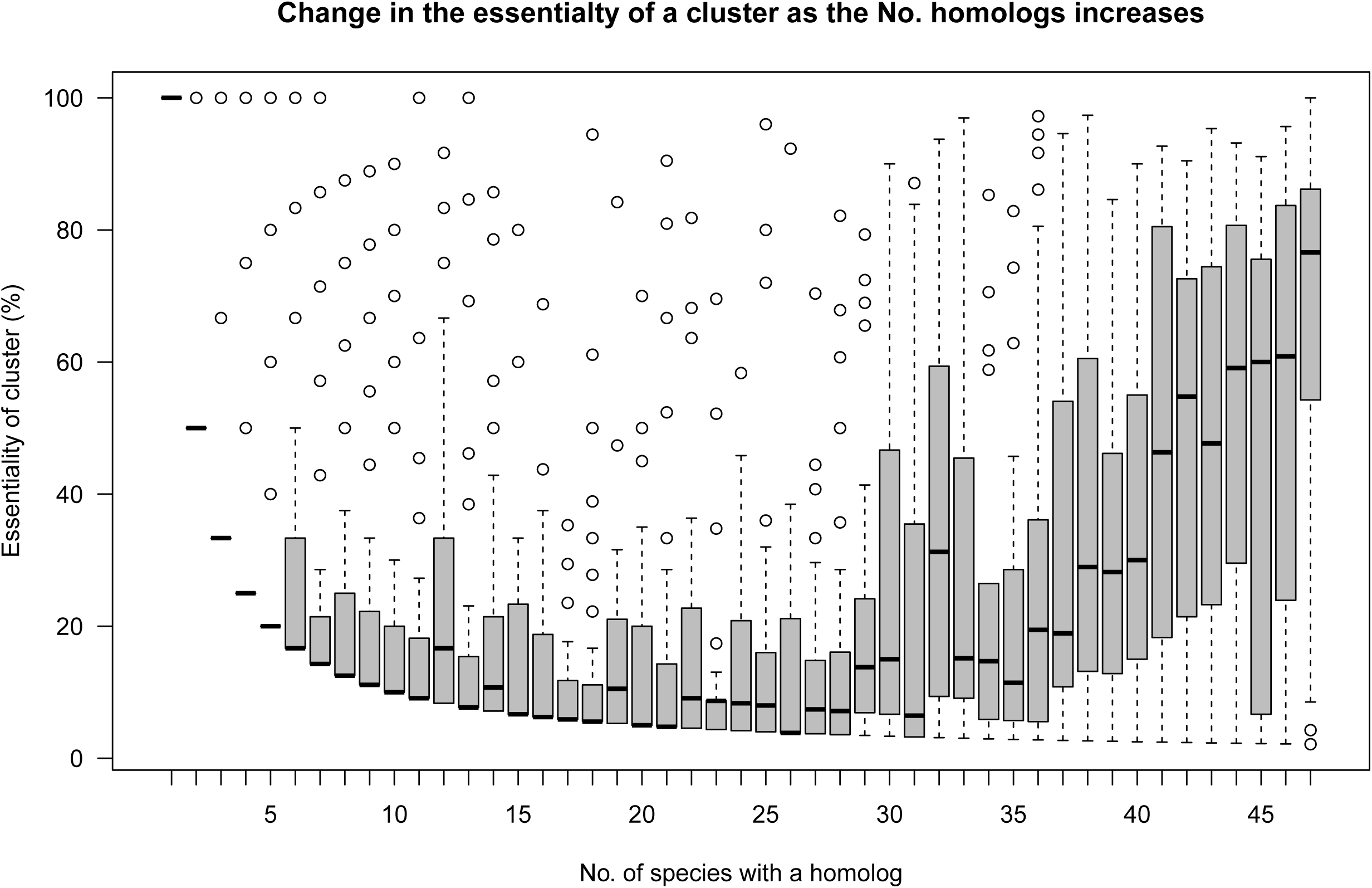
Changes in the essentiality of a cluster of homologs containing an essential gene. By studying the conservation of essential genes we have found a bi/modal trend in essentiality when the number of homologs increases.

## DISCUSSION

Within this study, out of 47 species we found 92 genes with a homolog present in every species analysed, and 127 if we assumed that a gene could not be found in a species due to annotation error or search problems. Comparison between our dataset and those generated by Charlebois and Doolittle (17), Koonin (15) and Lan et al., (18) show a high degree of overlap. While not all species shared an exact homolog, the genes deemed universal by Charlebois and Doolittle (17) were present in all our species, along with 90% of the genes identified by Lan et al., (18). This list of universally conserved genes is dominated by COG category J, translation and biosynthesis, which is in line with previous analyses (15,17). The remaining COG categories have very few genes shared between all species. This is most evident in the genes regarding transcription (COG category K) and the genes involved in metabolism. Transcription initiation is a vital cellular processes, yet there are only four genes that are conserved across all species: the Holliday junction resolvase *ruvX*, transcription termination/anti-termination protein *nusA*, and the two DNA directed RNA polymerase subunits A & B, *rpoA* and *rpoB*. These four proteins are responsible for the process of transcription, and functions relating to its termination, but there are no universal transcription initiation proteins.

The lack of universal transcription factors is interesting, as is the general lack of essentiality within the class. Of the 10829 genes in the database that belong to COG category K, only 712 are essential. While this is not the lowest percentage of essentiality for a COG at 6.6%, compared to the other COGs in its cluster (J has 40.4% essentiality and L has 19.9%), it is a significant change. By contrast, the transcription genes are far more diverse than the genes relating to translation and DNA replication and repair, as transcription in bacteria is a highly diverse process. Many bacteria rely on a vast array of different, niche specific transcription factors, along with other factors such as supercoiling DNA and nucleoid associated proteins (41–43), all of which are far more species specific than the fundamental DNA repair & replication and translation machinery.

One of the problems we have in our analysis is that the phyla belonging to the Proteobacteria comprised 33 of 47 species in our analysis. Thus, a bias towards this phyla’s genetic predispositions is inevitable. This is not just an issue specific to this study, but found across microbiology in general. A review of the GenBank entries regarding sequenced bacterial genomes in 2015 found that just six bacterial phyla comprise 95% of all sequenced bacterial genomes, and 46% of the total sequences were from Proteobacteria (44). Of the remaining phyla, in order of number of genomes sequenced, were the Firmicutes (31%), Actinobacteria (13%), Bacteroidetes (3%), Spirochaetes (2%), Cyanobacteria (1%), then all other phyla (5%). Therefore, while our ratios are slightly different, this study did analyse data that is generally representative of the overall state of sequenced bacteria. Because of the bias towards Proteobacteria, there is a second implicit bias towards Gram-negative bacteria. This could be due to the fact that Gram-negative bacteria are intrinsically more receptive to transformation due to their lack of peptidoglycan cell wall, thus essentiality studies on them are easier to perform. The thick peptidoglycan layer acts as a natural barrier for transformation and makes methods such as electroporation less effective, though polyethethylene glycol has been shown to be effective (45).

Accounting for experimental noise is a key issue with this analysis. Analysing transposon data for essentiality is inherently prone to many confounding factors, and eventually based on statistical probabilities instead of empirical observation (25,46). Because of this, by combining multiple different results, each utilising variations in methodology, such as choice of transposon, transformation method, growth condition(s) and analysis pipeline, this noise will propagate throughout this study. The case of *B. thuringiensis* shows that there is a level of noise generated by the described issues in reporting and annotation of essential genes. As such, while it may be feasible or useful to conclude that there is a strong chance that genes with a percentage essentiality of 89% or higher can be assumed to be universally essential; there are certain to be individual cases that do not conform. Being strict we only find *DnaA* as universally essential, and when adopting the 89% cutoff we find 26 genes which are essential across all organisms analysed here, and their function is dominated by DNA replication & translation machinery (see Table 1).

However, we must be careful ascribing the lack of 100% essentiality to experimental errors. This is exemplified by the S-adenosylmethionine gene *metX*. This gene is found in all species, and is a major component of the methylation systems of both DNA, RNA and proteins (47). It is essential in 89% of species, but is non-essential in many of the larger bacteria. For example, *A. tumefaciens* contains a secondary adenine methyltransferase known as *CcrM*, which is essential (48). This gene performs a highly similar function to *metX*, and could explain the loss of essentiality of the latter gene. This inherent probability that there are paralogs or moonlighting functions for highly conserved genes shows why it is probably not feasible to ascribe a hard delineation to the number of essential genes within the population. As it is the phenotype that is essential, and multiple genotypes can allow for this, it may be more accurate to attempt to define the essential phenotypes needed for life and work out the essentialities relating to them. Alternatively, we can treat the percentage of essentiality of a gene as a confidence level that it is truly essential across a disparate population of bacteria.

This phenomenon is shown clearly when looking at the absence of certain genes from the conserved list. A noteworthy absentee is the Sigma 70, which is present in 38 species (80%) and essential only in 68% of the species it is found. A member of the sigma 70 is a ubiquitous requirement for bacteria, as its role in housekeeping transcription is supposedly vital (49). Here, we find that while the canonical Sigma 70 is well conserved, a large number of species contain non-homologous members of the sigma 70 protein family. Along with its relatively low level of essentiality across the dataset, this implies that despite its fundamental role in housekeeping transcription, there must be other members of the sigma family that can provide the same function, at least in *in vitro* conditions.

The almost universal preservation of many NE genes shows that high level conservation of a gene is not dependant on it being essential to cellular survival. However, this could be a good example of the role of genetic redundancy and the divide between an essential function vs an essential gene. Some of these genes could truly be NE in some organisms, however they may also contribute to an essential function. Of the 57 near universally conserved but rarely E genes, 33% are involved in DNA replication & repair Having multiple redundant systems to repair DNA damage could impart a strong survival advantage *in situ*, which in turn is not needed for highly controlled laboratory conditions. For example, the *UvrABC* system is nearly universally conserved, and nearly universally NE. However, DNA damage via UV irradiation is controlled against *in vitro*, thus the function becomes NE in the conditions it was tested under.

The identification of these proteins, and those like them, could be highly useful in regards to genome engineering. Even the most ambitious genome reduction projects have had to retain certain NE genes (2). Knowing which are highly conserved, and thus most likely to have a strong fitness impact on the cell under certain conditions, could help guide rational design of new genomes beyond the simple dichotomy of preserving E genes and dispensing with as many NE genes as possible.

As the complexity of the genome increases, new reaction pathways are added, and thus become integrated into pre-existing circuits. This can make some genes become NE, as new genes bring with them redundancies for pre-existing ones, but it can also make pre-existing genes E. This can be due to the fact the substrate they produce is now vital to the proper functioning of a new pathway, or is somehow involved in its regulation, thus leading to an increase in the E genes. A good example of this is found within the lipid metabolism pathways. Organisms in the ‘minimal’ or ‘small’ genome category have a reduced ability to synthesise their own lipids (50). The 1-deoxy-D-xylulose 5-phosphate reductoisomerase gene *dxr* is the first step in the isopentenyl diphosphate biosynthesis pathway. It is present in 41 of the 47 species and is essential in 38, absent in the three mollicute species analysed here, the two streptococci and *L. crescens*, all of which are in the ‘minimal’ or ‘small’ genome categories. This could explain the small correlation between genome size and the number of E genes (Pearson’s Product-Moment correlation of 0.28) that we find.

The reason perhaps why this correlation is not very high could be due to the fact that as the number of genes within a genome increases, more and more pathways are added and thus pathways can be replaced, increasing genetic redundancy. A good example of this can be seen in the genes related to carbohydrate metabolism in Suppl. Figure 13. Among the highly conserved genes, there is a clear tendency for genes to be essential in the smaller genomes and non-essential in the larger ones. This is also seen in the genes involved in translation (COG category J, Suppl. Figure 12), though to a much smaller extent.

The bi-modal trend of the essentiality of the genes changing with the number of homologs present fits well with our hypothesis of increasing complexity and gene utilisation. Genes only found in a single organisms are likely to be there as a response to some form of environmental stress specific to the niche that bacteria inhabits. This trait is especially true in pathogenic bacteria, which tend to evolve similar orphan genes when dealing with similar pathogenic niches. These genes are rarely if ever found in non-pathogenic species, even within the same genus, implying that there is a strong evolutionary pressure stemming from their niche which these genes help alleviate (51). However, as genes become present in more and more species, this implies that their functionality becomes useful to a wider range of niches. The more widely conserved a gene is, the more likely to encode for a protein that assists in a general or housekeeping function instead of a niche specific one, thus the more likely it is to have some level of genetic redundancy. Finally, genes that are nearly universally conserved, by definition must play a role in a fundamental cell process. While there will be some level of redundancy in its functionality, the fact that none of the species analysed has replaced the original protein with a redundant or modified one implies that the function it provides is still vital for cellular function at a fundamental level.

Finally, it is probable that the potentially unclassifiable variation in bacteria (11), coupled with the abilities of genes to moonlight to other functions (52–54), especially in pathogenic bacteria (55– 57) could explain the lack of commonality in gene conservation and essentiality. This potential for bacteria to employ similar genes to fill multiple roles allows for a level of redundancy within key processes, enabling bacteria to specialise their function, and perfectly suit their niche (58).

Therefore, while certain cornerstone functions of life seem conserved across the Bacterial Domain, bacteria have evolved many redundancies and replicate systems to allow for differentiation. Even the ribosome, one of the most conserved structures across all domains of life, cannot be fully re-created from the conserved proteins, and many of those proteins are NE in many species. This lack of consensus between E genes and E functions highlights the need to understand the functionality of all genes, and their cumulative role in cell processes, if we are to fully understand what components truly are and are not essential for all bacterial life.

## AVAILABILITY

Genome Maps is an open source collaborative initiative available in the GitHub repository (https://github.com/compbio-bigdata-viz/genome-maps)

## SUPPLEMENTARY DATA

Supplementary Data will be available online.

## ACKNOWLEDGEMENT

N/A

## FUNDING

This project has received funding from the European Union’s Horizon 2020 research and innovation programme under grant agreement 634942 (MycoSynVac) and was also financed by the European Research Council (ERC) under the European Union’s Horizon 2020 research and innovation programme, under grant agreement 670216 (MYCOCHASSIS). We also acknowledge support of the Spanish Ministry of Economy, Industry and Competitiveness (MEIC) to the EMBL partnership, the Centro de Excelencia Severo Ochoa and the CERCA Programme/Generalitat de Catalunya.

## CONFLICT OF INTEREST

None declared.

